# 3D reconstruction of ultra-high resolution neurotransmitter receptor atlases in human and non-human primate brains

**DOI:** 10.1101/2022.11.18.517039

**Authors:** Thomas Funck, Konrad Wagstyl, Claude Lepage, Mona Omidyeganeh, Paule-Joanne Toussaint, Katrin Amunts, Alexander Thiel, Nicola Palomero-Gallagher, Alan C. Evans

## Abstract

Quantitative maps of neurotransmitter receptor densities are important tools for characterising the molecular organisation of the brain and key for understanding normal and pathologic brain function and behaviour. We describe a novel method for reconstructing 3-dimensional cortical maps for data sets consisting of multiple different types of 2-dimensional post-mortem histological sections, including autoradiographs acquired with different ligands, cell body and myelin stained sections, and which can be applied to data originating from different species. The accuracy of the reconstruction was quantified by calculating the Dice score between the reconstructed volumes versus their reference anatomic volume. The average Dice score was 0.91. We were therefore able to create atlases with multiple accurately reconstructed receptor maps for human and macaque brains as a proof-of-principle. Future application of our pipeline will allow for the creation of the first ever set of ultra-high resolution 3D atlases composed of 20 different maps of neurotransmitter binding sites in 3 complete human brains and in 4 hemispheres of 3 different macaque brains.

## 1. Introduction

### 1.1. Neurotransmitter receptor atlas

Three dimensional (3D) digital brain atlases are essential to the analysis of brain structure-function relationships, but, as of yet, high resolution atlases of human neurotransmitter receptor maps have not been available. The lack of neurotransmitter receptor maps is particularly unfortunate given that neurotransmitters and their corresponding receptors underpin all synaptic transmission and hence all information processing in the brain.

While atlases based on in vivo neurotransmitter receptor imaging using PET maps have recently been created^1,2^, PET suffers from relatively poor spatial resolution compared to the scale of regional and laminar distribution of neurotransmitter receptors. This lack of resolution makes it difficult, and sometimes impossible, to resolve important biological features of the cortex such as both areal and especially laminar patterns of receptor or cell-body distributions.

We have developed a pipeline for creating 3D atlases of human and macaque brains, each of which is composed of maps from 2D sections, including receptor autoradiographs, cell body stained sections, and myelin stained sections. While the pipeline can be applied generically to virtually any post-mortem 2D sections where the cortical grey matter (GM) can be visualized, we here apply it primarily to a data set composed of autoradiographs with the goal of reconstructing 3D 20μm^3^ atlases of neurotransmitter receptor distributions in the cortex for both human and macaque brains. Future work will enable integration of subcortical regions to these atlases based on a parallel volumetric reconstruction workflow.

### 1.2 Data Acquisition and Challenges

The autoradiography data set collected by Karl Zilles over a period of almost two decades (for overviews see, Zilles et al.^3^; Palomero-Gallagher and Zilles^3,4^) is unique in the world and exceptionally rich, because it samples 20 different neurotransmitter receptor binding sites across both hemispheres of 3 human brains, and in macaque data, 15 different neurotransmitter receptor binding sites, cell body and myelin stained sections from 3 hemispheres from 3 macaque brains. Furthermore, MRI datasets are also available for the human brains. During the acquisition, the brain is extracted from the cranial cavity immediately following death, as receptor binding of autoradiographs requires processing of fresh tissue, cut into slabs (see Figure 1A-B) and shock frozen, then sectioned and stained. Until now, these data sets of 2D autoradiographs, cell body stains, and myelin stains had never been reconstructed into 3D because of the numerous and considerable challenges associated with data collection and processing.

**Figure 1.**
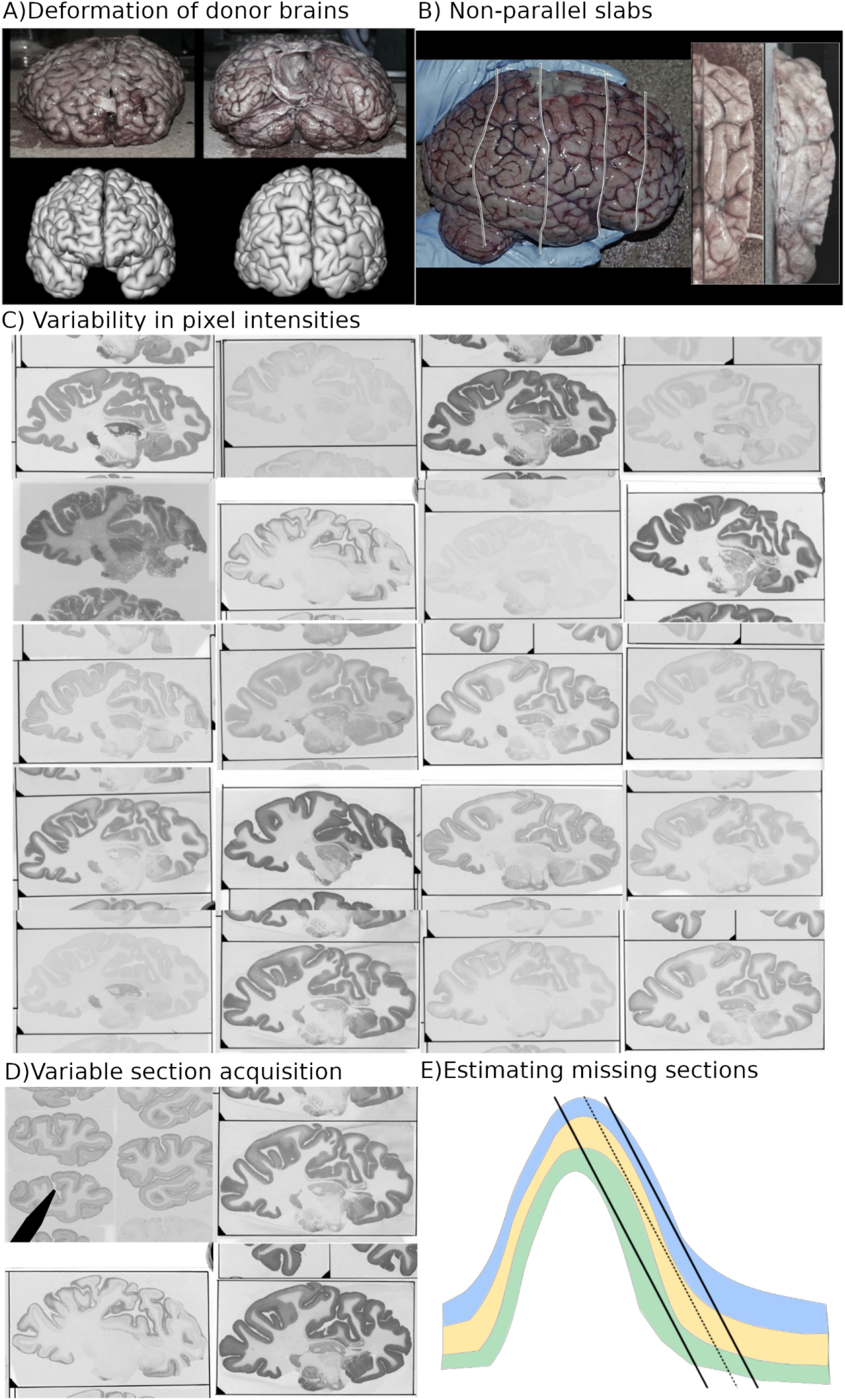
A) Donor brains were not chemically fixed and hence exhibited significant deformation. B) Tissue slabs were not cut parallel to one another and substantial gaps were present between acquired slabs. C) Exemplary autoradiographs from the human data set showing each of the 20 neurotransmitter receptor binding sites illustrating the substantial heterogeneity in pixel intensities between the autoradiographs. D) Acquired sections have multiple pieces of tissue and visual cues that must be removed prior to reconstruction. E) Histological sections acquired tangentially to the surface curvature of the cortex (the solid lines) will misrepresent the laminar distribution of neurons, neurotransmitter receptors, and other biological features. Estimating missing pixel intensities between such sections (the dotted line) by averaging the two sections would propagate this sectioning artefact. Therefore, some method is required to accurately estimate missing pixel intensities between acquired sections.

For simplicity of terminology, all 2D images acquired post-mortem after sectioning a brain are referred to as histological sections, where different types of histological sections may represent cell body density, myelin density, or the density of receptor binding sites. As our reconstruction pipeline aims to be agnostic to the method of acquisition, these images are treated equivalently. The information represented in these images is referred to generically as pixel intensities.

#### 3D non-linear warping

The donor brains were not chemically fixed prior to removal from the cranial cavity, since this would affect receptor binding properties^4^, resulting in significant non-linear warping (Figure 1.A). Moreover, not only is the brain as a whole subject to non-linear deformations, the individual slabs into which the hemisphere had to be cut in order to minimise artefacts during shock freezing are subject to their own deformations prior to freezing.

#### Non-parallel slabs

Prior to sectioning of the human donor brains into 20µm thin 2D sections, the fresh brains were cut into slabs of tissue of approximately 2-3cm along the coronal axis (Figure 1.B). These cuts were not perfectly parallel to one another. This created slabs that were not sectioned along uniform parallel planes and whose exact position in the brain is not obvious. Therefore the pipeline must both align the sections within a tissue slab to one another and then align the tissue slab to the reference structural volume.

The macaque brains were cut into two slabs prior to freezing, but were treated as a single slab for the purpose of reconstruction.

#### Variability in pixel intensity distribution from different histological acquisitions

One of the most important obstacles for 3D reconstruction of the present data was the degree of variability in the pixel intensity distributions of histological sections. This is illustrated in Figure 1.C, where the autoradiographs from the human brain reveal very different receptor distributions and overall image contrasts, and the same applies to autoradiographs obtained from the macaque brain. An automated pipeline must perform equally well for the various pixel intensity distributions.

Aligning the 3D histological volume of each tissue slab directly to the reference structural volume is therefore challenging because of the relatively sparse sampling of the histological sections and their heterogeneous pixel intensities, both within and between sections.

#### Variability in autoradiograph acquisition protocol

The raw histological sections, primarily the autoradiographs from the human data set, were acquired with multiple brain sections and non-brain visual cues (fiducial frames, arrows, circles, and triangles; Figure 1.D) on a single photographic film. This requires an automated cropping step to identify the target piece of brain tissue from each digitised section and remove extraneous pieces of brain tissue and non-tissue visual cues from the image.

#### Estimation of pixel intensities for missing histological sections

When performing serial acquisition of multiple types of histological sections, e.g., receptor, cell body, and white matter (WM) density, there are necessarily significant gaps between acquired sections of a given type. This problem is compounded by the loss of sections due to mechanical processing. The sparse sampling of histological sections necessitates a method for estimating the missing sections for a particular type of histological section. This is particularly difficult given that the cutting angle of the section will often not be perpendicular to the curvature of the cortex (Figure 1.E). That is, to the extent a section is cut tangentially to the curvature of the cortex, it may obscure the laminar distribution of biological features, such as cell body or receptor density, across the cortical depth. Hence an interpolation method is required for estimating missing histological sections that will not propagate artefacts resulting from the cutting angle of the sections.

### 1.3. 3D reconstruction from 2D post-mortem brain sections

Many reconstruction algorithms have been proposed (see Dubois^5^ and Pichat et al^6^ for reviews), but none of these conventional methods are adapted to solve all of the challenges presented in 1.2.

Semi-automated 2D reconstruction methods have been proposed using fiducial markers implanted in the brain prior to sectioning^7,8,9^ and using block-face imaging ^10,11,12^. However, neither fiducial markers nor block-face images were used in the acquisition of our data, so these techniques are not applicable in the present case.

Another semi-automated approach to 2D image reconstruction was to manually identify anatomic landmarks on adjacent sections^13,14^. While such methods could in principle work for the initial alignment of our data, they are labour intensive and dependent on rater subjectivity.

Automated reconstruction can be performed with only the 2D sections themselves using principal-axes transforms^15^, intensity or frequency-based cross-correlation^16,17^, sum of squared error^18^, discrepancy matching optical flow^18–20^, or edge-based point matching^18–20^. The main drawback of these methods is that misalignment errors will be propagated to all subsequently coregistered sections^18,19^.

Methods have been developed to perform more robust alignment between sections and to maximise the smoothness of the 3D reconstruction^21–24^. While these methods have been shown to perform well in their respective applications, they are unlikely to perform as accurately on the present data set. This is because of the variable intensity distributions in the autoradiographic sections and the considerable anatomic variability stemming from the gaps between acquired sections.

A recent and particularly innovative approach to 2D alignment of histological and corresponding MRI sections involved using Bayesian methods to simultaneously align the histological section and the MRI section while transforming the pixel intensities of the former to resemble the latter^25,26^.

Finally, an iterative strategy for reconstructing 2D sections using an accompanying structural brain image was proposed by Malandain et al^27^ and later adapted by Yang^28^ and Amunts et al^29^. This scheme iterates between two steps at progressively higher spatial resolutions. First the reference structural brain image is aligned in 3D to a stack of 2D sections and then the 2D sections are linearly aligned in 2D to the structural brain image. By beginning at a coarse spatial resolution and progressively refining the resolution, the pipeline converges to an accurate alignment between the 2D sections and the 3D reference structural volume.

### 1.4 Reconstruction pipeline overview

To address the unique challenges presented by the 3D reconstruction of our histological data set, we have developed a mostly, though not fully, automated pipeline that expands on the pipeline presented by Malandain et al^27^ and Amunts et al.^29^ by creating a multi-resolution framework of non-linear alignment that is compatible with sparsely sampled data and supplementing a surface-based interpolation algorithm for estimating missing pixel intensities. Furthermore, as neurotransmitter receptor binding is evidenced in gray matter, registration is simplified to mono-modal alignment of gray matter intensities.

Briefly, the reconstruction pipeline (Figure 2) is composed of 4 major processing stages.

1. Automated cropping of the sections to isolate the target piece of tissue.
2. An initial histological volume is created by rigid 2D inter-section alignment.
3. An iterative multi-resolution alignment scheme that alternates between 3D volumetric followed by 2D section-wise alignment of the histological sections to the reference structural brain image (e.g., donor’s T1w MRI). The alignment between the histological volume and the reference structural volume is performed using binary grey matter (GM) volumes derived from each of these data sets, respectively. The problem of aligning a histological volume composed of heterogeneous pixel intensities to a reference volume with an entirely different pixel intensity distribution is thus simplified to mono-modal alignment between GM volumes.
4. Morphologically informed surface-based interpolation is used to estimate missing pixel intensities for locations where a type of histological section was not acquired.

**Figure 2.**
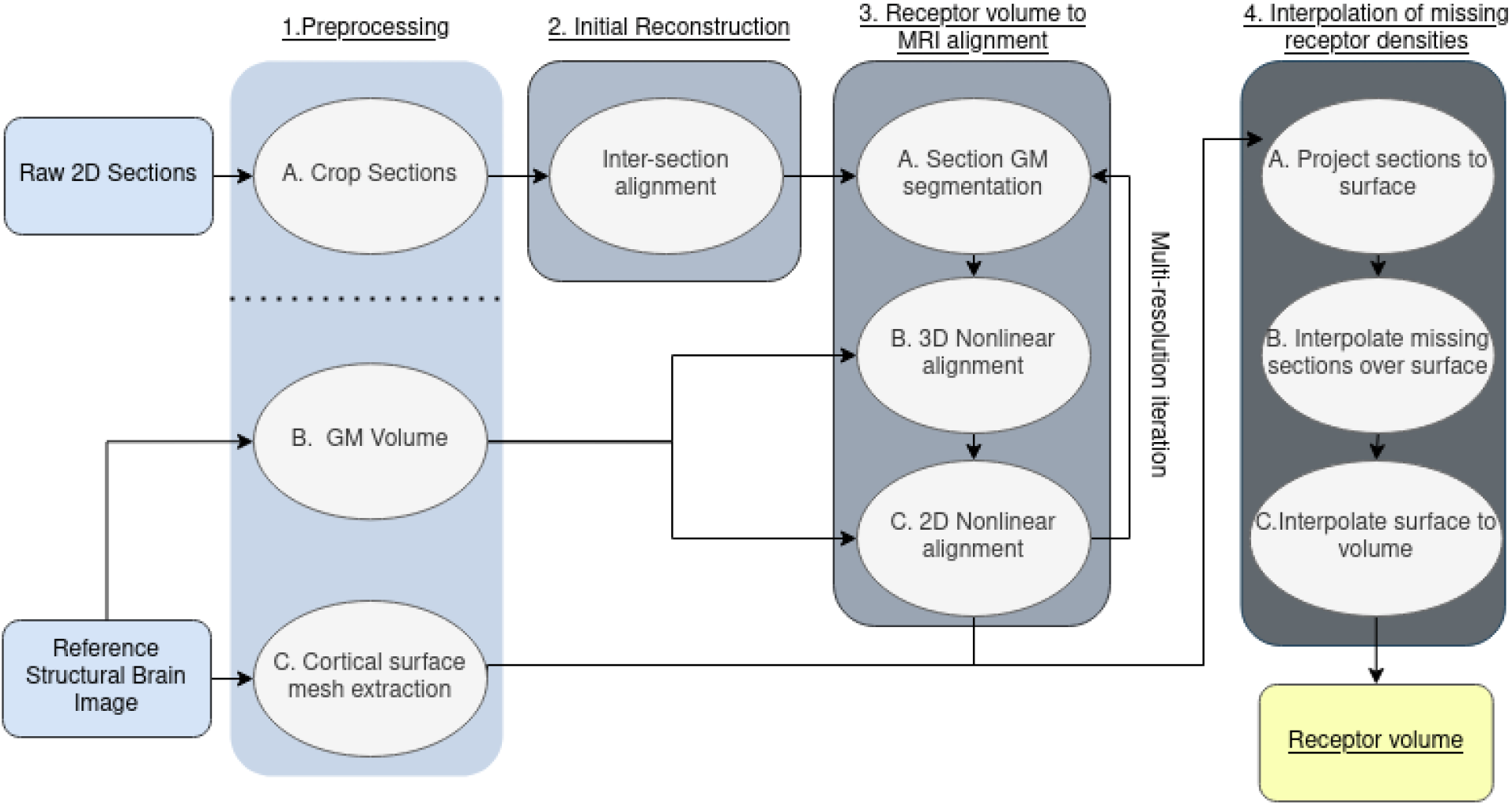
Overview of the reconstruction pipeline. The pipeline contains 4 major processing stages: 1) semi-automated cropping of the sections, 2) inter-section 2D alignment, 3) iterative multi-resolution 3D volumetric followed by 2D section-wise alignment of section to the reference structural brain image, 4) surface-based interpolation of receptor binding densities.

We validated the reconstructed densities using the pixel intensities, either binding site densities or cell body packing density, in the original 2D sections.

## 2. Results

The alignment of the histological sections acquired from the human donor is seen in Figure 4 (left) and the macaque in Figure 4 (right). The non-linearly aligned 2D histological sections are shown in a sagittal and axial view with WM and GM cortical surfaces overlaid on top. The alignment is generally accurate but requires further improvement in some regions such as the temporal lobe.The reconstruction performed similarly between the human reconstruction with a donor MRI and the macaque reconstruction using a reference template brain.

The accuracy of the human reconstruction was quantified by calculating the Dice score between aligned histological sections and corresponding sections from the structural reference volume. The average accuracy of the alignment was 0.91±0.1. The largest source of error was at the ends of the tissue slabs were incomplete tissue sections were acquired (Figure 5).

The reconstruction pipeline was able to create receptor binding volumes for all 20 types of binding sites for the human brain (Figure 6.A). Additionally, the pipeline was able to produce volumes of GABAA/BZ binding sites, cell body, and myelin density volumes in the macaque brain (Figure 6.B). The pipeline is therefore able to accurately align sections from a non-human primate using a reference template brain MRI instead of the donor MRI.

The reconstruction pipeline was validated to quantify the amount of error induced in the pixel or voxel intensities due to resampling and interpolation. The accuracy of the measured pixel intensities was calculated after 2D nonlinear alignment (A.3.2) and after full 3D reconstruction (A.4.4). The accuracy in both stages is ~98%, but there is more variance in the accuracy after applying the 3D nonlinear transform, especially for ROIs with smaller volumes (Figure 7). The standard deviation of the accuracy was 2.2% after 2D alignment and 3.8% after 3D reconstruction. The variability in accuracy of the pixel intensities in the reconstructed volume appears to improve for ROIs of at least 100-200μm^2^.

A surface-based interpolation algorithm was used to estimate missing pixel intensities between acquired autoradiographs. The surface-based interpolation was validated by applying it within randomly selected patches of vertices within acquired autoradiograph sections. For each ligand, the overall correlation between true and interpolated pixel intensities was r^2^=0.98±0.001, with only a few outlier values where the interpolation performed poorly (Figure 8). The distances between the vertices with known pixel intensities and the vertices to be estimated spanned 0.05-1.2mm.

## 3. Discussion

### Summary

We have created a semi-automated pipeline that can successfully reconstruct 3D volumes from multiple histological sections, specifically volumes of binding densities for 20 different neurotransmitter receptor binding sites from a human donor brain and volumes of GABAA/BZ, cell body, and myelin density from a macaque brain. To recapitulate briefly, the pipeline automatically cropped the histological sections to isolate target brain tissue. The cropped histological sections were then aligned to one another using rigid transformations to create an initial 3D histological volume. For each tissue slab, respectively, an iterative multiresolution scheme was then used to align the histological volume to the reference structural brain image. Finally, for each kind of histological section, a surface-based interpolation algorithm was used to create volumetric maps that represent the distribution of the biological feature measured by each respective histological acquisition method.

Visual and quantitative validation of the quality of the alignment of the histological sections indicated that the reconstruction pipeline can accurately recover the 3D anatomy of the reference volume from the 2D histological sections (see Figure 4). The average Dice score of the aligned histological sections to the corresponding sections from the reference structural volume was 0.91±0.1. However the accuracy of the alignment was much lower at the ends of the slabs (see Figure 5) because the sectioning at the edges of the tissue slabs produced fragmented sections with only a small portion of the brain (see Figure 5.B). As it was unclear which histological sections the pipeline would be able to correctly align, all sections were included in the reconstruction. However, when producing reconstructed volumes for public use, these sections will either have to be removed or, if possible, manually aligned. The sharp drop in Dice scores will provide an automated way of detecting at which point the sections from a tissue slab can no longer be used in the reconstruction.

The quantitative validation of the alignment will also allow us to create “error maps” that indicate where the reconstruction has performed less well and where users need to exercise caution when using the data.

The quantitative validation of the pixel intensities shows that the reconstruction pipeline preserves pixel intensities with 98% accuracy but with increased variance after 3D alignment of the histological sections, especially for smaller ROI (Figure 7). The increased variance after 3D alignment is inevitable due to in-plane and out-of-plane interpolation errors induced by warping the 2D sections in 3D space. However the overall accuracy for a given size of ROI will improve when the reconstruction is performed at the target resolution of 20μm^3^ because there will be no in-plane interpolation error due to downsampling.

The quantitative validation of the surface-based interpolation demonstrates that estimated receptor intensities are highly correlated to the true pixel intensities, r^2^=0.98±0.001 error (Figure 8), demonstrating that the surface-based interpolation algorithm is able to estimate accurate pixel intensities over the surface of the cortex. The surface-based interpolation algorithm was applied to estimate vertices from 0.05-1.2mm away from vertices with known pixel intensities. In the human data, the average sampling between histological sections of the same type was approximately 570μm.

The results also demonstrate that the reconstruction pipeline is not limited to human brains but can be applied to macaque brains as well. Moreover, the reconstruction of the macaque brain was performed without a donor MRI, using the MEBRAINS stereotaxic atlas instead. This is significant because large amounts of 2D brain sectioning data have been acquired without corresponding 3D structural imaging, such as MRI. Our method can therefore reconstruct data sets of 2D sections acquired with histology, autoradiography, polarised light imagery, etc, but where no corresponding 3D structural image has been acquired. An important caveat to note is that macaque brain morphology is much simpler than that of humans. Therefore, while it may not be necessary to have a corresponding 3D structural image when reconstructing sections from animals with relatively simple cortical morphology, it may likely be necessary to reconstruct sections acquired from human brains.

### Comparison With Existing Methods

A similar scheme was used by both Malandain et al.^27^ and Amunts et al.^29^ to reconstruct histological volumes in 3D. Generally speaking, this scheme iterates between two steps: first the donor MRI is aligned in 3D to a stack of histological sections and then, in the second step, the histological sections are linearly aligned in 2D to the transformed MRI. Amunts et al.^29^ improved on this approach by using a multi-resolution nonlinear warping schema.

Unfortunately, their approach cannot be directly used with sparsely sampled sections with heterogeneous pixel intensity distributions. An important novel method we implemented for overcoming the problems with this sort of data was recalculating a continuous GM volume between acquired sections at each resolution. This means that at each resolution we can create a smoother, continuous 3D representation of the histological sections which permits more accurate spatial alignment to the reference structural volume. This approach allows for the reconstruction of even very sparsely sampled histological sections as in the macaque data where sections were acquired in chunks approximately 2.6mm from one another.

A fundamental difference between our approach and existing methods is the introduction of morphologically informed estimation of missing pixel intensities using cortical surface-based interpolation. The interpolation is performed over the cortical surface to account for confounds that result when the cortex is sectioned at an angle that is not perpendicular to the curvature of the cortex. These artefacts may result in layers of pixel intensities within the cortex, e.g. cytoarchitectonic layers, being obscured within the 2D section. Simply interpolating between two sections through Euclidean space, e.g., Euclidean distance-weighted linear interpolation between two sections, would propagate this sectioning artefact to the estimated pixel values.

The impact of sectioning artefacts is limited by projecting pixel intensities onto the cortical surface meshes at equidistant depths across the cortex and then performing linear interpolation over the geodesic distances of the cortical manifold. The advantage of this approach is that cortical surfaces and geodesic distances better represent the actual morphology of the cortex than Euclidean distances.

Many methods have been proposed to create smooth and continuous 3D volumes from 2D sections without the use of external references^22–24^, such as fiduciary markers, block face images, or structural reference volumes. In principle these methods would be helpful for creating the initial reconstruction in Section A.2. However, to our knowledge, these methods cannot be directly applied here again because of the sparse sampling of histological sections with heterogeneous pixel intensities. Fortunately, because the multiresolution schema described in Section A.3 begins at a very low spatial resolution of 4mm^3^, it only requires that the histological sections be grossly aligned to start with.

Another promising approach has been proposed by Iglesias et al^25^. Their method uses Bayesian inference to estimate both physical transformations to the sections and the transformation of pixel intensities to align sections acquired with different intensity distributions. Our method, by contrast, uses two different approaches. When aligning 2D sections directly, the use of an information theoretic similarity metric proved sufficient for accurate alignment. However, when aligning a slab of tissue composed of sections with many different intensity distributions to a reference structural volume, such as a T1w MRI, we simplified the problem of multi-modal alignment by transforming all sections to binary GM image. This simplification makes it possible to align a volume composed of many tissue types to a reference structural volume.

### Limitations

#### Interpolation of Missing Sections

The sequential acquisition of different types of histological sections means that there is a minimum gap between acquired sections of the same type of at least 400μm for the human data and 1mm for macaque data. This is a fundamental limitation of the data set that cannot be improved on with the data at hand. An interpolation algorithm was devised to provide an estimate of the missing pixel intensities based on the nearest available sections. The surface-based interpolation scheme assumes that pixel intensities measured from histological sections change linearly between the acquired sections and the missing section. This is not strictly biologically valid because there may be sharp boundaries between, say, cytoarchitectonic areas^3,27^ which would be obscured by our interpolation method. It does not appear possible to devise an interpolation method that could reproduce such sharp regional boundaries without additional anatomic information.

#### Quantitative accuracy of autoradiography

An important underlying assumption in the use of autoradiography is that post-mortem in vitro radioligand binding reflects in vivo neurotransmitter receptor density. The available evidence suggests that prolonged freezing of brain tissue does not affect receptor binding sites^30–33^. The quantitative accuracy of autoradiography could, however, be affected by the delay elapsed between the donor’s time of death and the freezing of the brain. For example, NMDA, GABA, muscarinic M_1_, D2, and 5-HT_2_ receptor binding sites were stable for up to ~75 hours post-mortem^32,34–36^. Other neurotransmitter receptor densities increased post-mortem before freezing, e.g., D_1_ and 5-HT_1A_ receptor binding sites^37^. Most problematically for the present work, the benzodiazepine receptor binding sites increased by 150% within a 48h post-mortem delay^30^. However, given the post-mortem delay of less than 24 hours of the brain used for the present study, we can assume that the receptor densities encoded by the ensuing receptor volume reflect in vivo density.

#### Segmentation of histological sections

A potential limitation of our approach is that it relies on the ability to estimate an accurate GM image from a 2D section. It is possible that there are imaging modalities where this estimation of cortical GM may be more challenging. However, the network used in this work is trained using only synthetic data which can, in principle, be extended to include different tissue contrasts, more tissue classes, and additional imaging artefacts ^38^. This means that if there is a particular type of histological section that is not well segmented by the current network, the synthetic data set can be augmented to reflect the particularities of the problematic sections and hence improve the segmentation of these sections.

## Conclusion

We have created an image processing pipeline for reconstructing 2D histological sections into 3D volumes. The results here serve as a proof-of-principle that the pipeline can reconstruct the 2D autoradiographs accurately at high resolution. The work presented here will allow for the creation of an unparalleled, and publicly available, data set of 20 receptor binding site volumes at 20μm for 3 human brains and 3 hemispheres from macaque brains.

## 4. Material and Methods

### 2.1 Data Acquisition

#### 2.1.1 Human

In the present study, the brain of a single 78 year old male donor, obtained through the body donor program of the Department of Anatomy of the Heinrich Heine University Düsseldorf, was reconstructed as proof of principle. The donor died from non-neurological causes and had no neurological or neuropsychiatric clinical history. The subject had given written consent before death and/or had been included in the body donor program of the Department of Anatomy, University of Düsseldorf, Germany (three brains). An in situ post mortem T1w MRI was acquired on a Siemens Magnetom Sonata scanner with an MPRAGE acquisition protocol (2.2s TR, 1.2s TE, 15° flip angle).

The human brain was divided into left and right hemispheres, then cut into slabs of tissue of approximately 2-3 cm to facilitate the even freezing of the brain tissue (Figure 1.C). Each slab was shock frozen between -40 and -50 C in N-methylbutane with a post mortem delay of less than 24 hours in the case of the human brain.

Slabs were sectioned at -20°C into 20 μm thick sections of brain tissue with a large scale cryostat microtome and thaw-mounted on gelatin-coated glass slides. No blockface images were acquired. Sections were freeze-dried overnight prior to incubation. Sections were first preincubated for rehydration and to eliminate any endogenous substances that may bind to the target receptor.

Brain sections were then radiolabelled in one of two ways. For sections imaged for total binding of the ligand to the target receptor binding site, sections were incubated in a solution containing the titrated radiolabeled ligand. Alternatively, a subset of sections were imaged for non-specific binding by incubating the sections in a solution containing the radioligand as well as an unlabeled displacer. The established binding protocols used here (described in detail in Table 1 of Palomero-Gallagher and Zilles^4^) resulted in a non-specific binding which was less than 5% of the total binding. Thus, we consider total binding to be equivalent to specific binding^4^.

**Table 1:** List of the 20 radioligands and associated neurotransmitter receptor binding site.

| Radioligand | Receptor | Transmitter Family |
| --- | --- | --- |
| AMPA | AMPA | Glutamate |
| Kainate | Kainate | Glutamate |
| MK-801 | NMDA | Glutamate |
| LY 341,495 | mGluR2/3 | Glutamate |
| Muscimol (agonist) | GABA-A | GABA |
| SR95531 (antagonist) | GABA-A | GABA |
| CGP 54626 | GABA-B | GABA |
| Flumazenil | GABA-A associated Benzodiazepine binding site | GABA |
| Pirenzepine | Muscarinic M<sub>1</sub> | Acetylcholine |
| Oxotremorine-M (agonist) | Muscarinic M<sub>2</sub> | Acetylcholine |
| AF-DX384 (antagonist) | Muscarinic M<sub>2</sub> | Acetylcholine |
| 4-DAMP | Muscarinic M<sub>3</sub> | Acetylcholine |
| Epibatidine | Nicotinic $\alpha_4\beta_2$ | Acetylcholine |
| Prazosin | $\alpha_1$ | Noradrenalin |
| UK-14,304 (agonist) | $\alpha_2$ | Noradrenalin |
| RX 821002 (antagonist) | $\alpha_2$ | Noradrenalin |
| 8-OH-DPAT | 5-HT<sub>1A</sub> | Serotonin |
| Ketanserin | 5-HT<sub>2</sub> | Serotonin |
| SCH 23390 | D<sub>1</sub> | Dopamine |
| DPCPX | Adenosine 1 | Adenosine |

The labelled receptor binding sites encompassed the classical neurotransmitter systems glutamate, GABA, acetylcholine, dopamine, serotonin, and noradrenaline, as well as the neuromodulator adenosine (see Table. 1 for a detailed list). Series of sequentially acquired sections were incubated with a specific radioligand or processed for the visualization of cell bodies^39^ or of myelinated fibers^40^ such that there were at least 23 sections of brain tissue between any two sections incubated with the same radioligand. Lastly, the sections were rinsed to remove excess radioligand and stop additional binding, then air dried at room temperature. See Palomero-Gallagher and Zilles^4^, for a detailed description of binding protocols.

Radioactively labelled sections were co-exposed with plastic titrated standards (Microscales®, Amersham) with known radioactivity concentrations against β-radioation sensitive films (Hyperfilm, Amersham, Braunschweig, Germany or Carestream Kodak BioMax MR film) for 4-18 weeks. The ensuing autoradiographs were digitised with a CCD-camera and the Axiovision (Zeiss, Germany) imaging and processing system.

The pixel intensities of the standards were plotted against their respective radioactivity concentrations, which had previously been calibrated to reflect receptor densities in fmol/mg protein based on Equation 1, and a calibration curve was fit to these points.

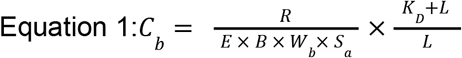

where *R* is the known radioactivity concentration of a standard, *E* is the efficiency of the scintillation detector, *B* is a constant for the amount of radioactivity decays per unit time (Ci/min), *W*_*b*_ is the protein weight of a standard (mg), and *S*_*a*_ is the specific activity of the ligand (Ci/mmol), *K*_*D*_ is the dissociation constant (nM) and *L* is the free concentration of ligand during incubation (nM). Finally, the grey value intensities in each pixel of an autoradiograph were transformed into binding site densities by interpolation from the calibration curve, thus resulting in linearized images (Palomero-Gallagher and Zilles^4^).

#### 2.1.2 Macaque

We acquired serial histological sections from one adult male macaque monkey brain (*Macaca fascicularis*; brain ID: 11530; 6 years of age; obtained from Covance Laboratories, Münster, Germany). The subject was euthanized with intravenous sodium pentobarbital and the brain was immediately extracted together with meninges and blood vessels to preserve cortical layer I. The procedures used in this study, in approval with the Institutional Animal Care and Use Committee, were carried out in accordance with the European and local Committees and complied with the European Communities Council Directive 2010/63/EU. No corresponding MRI was acquired for the macaque brain.

The brain was removed from the skull, and the brain stem and cerebellum were dissected off in close proximity to the cerebral peduncles. The brain was divided into 2 hemispheres by cutting the corpus callosum and was then cut in a rostral and a caudal block by cutting in the coronal plane of sectioning between the central and arcuate sulci. The unfixed tissue blocks were frozen in isopentane at −40 to −50 °C, and then stored in airtight plastic bags at −70 °C. Each block was then sectioned in the coronal plane using a cryostat microtome (CM 3050, Leica, Germany), obtaining sections of 20 μm thickness which were thaw-mounted on gelatine-coated slides and freeze-dried overnight.

Alternating sections were stained for cell bodies^39^ or myelin^40^, or processed for the visualization of neurotransmitter receptor binding sites. Specifically, tissue blocks were serially sectioned in such a way that groups of 25 sections (“repeats”) were collected throughout the slab, and 20 sections were discarded between repeats. Repeats consisted of a predetermined order of sections meant for the visualization of a specific receptor type or histological staining. Every 4th and 15th sections of a repeat were used for visualization of cell bodies, and every 9th and 20th sections for the myelin stain. Thus, the distance between two sections processed for the same histological type was 900 μm.

Quantitative in vitro receptor autoradiography was applied to label 15 different receptors for the transmitters glutamate (AMPA, kainate, NMDA), GABA (GABAA, GABAB, GABAA associated benzodiazepine [GABAA/BZ] binding sites), acetylcholine (muscarinic M1, M2, M3), noradrenaline (α1, α2), serotonin (5-HT1A, 5-HT2), dopamine (D1), and adenosine (A1) by incubating the sections in solutions of respective tritiated ligands according to protocols described in section 2.1.1.

As no MRI was acquired for the macaque brain, the MEBRAINS templatec brain was used as the anatomic template to which the autoradiograph sections were acquired.

### 2.2 Reconstruction Pipeline

In the first step of the pipeline (Figure 2.1), both the histological sections and the structural reference MRI volume must be preprocessed. For the histological sections, the goal of the preprocessing (see Supplementary Information A.1 for detailed explanation) is to isolate the target piece of brain tissue from each image and remove extraneous tissue and visual cues. For the reference donor brain, WM-GM and GM-pial surface meshes are extracted along with a binary cortical GM volume.

Once the preprocessing is complete, an initial reconstructed histological volume is created (Figure 2.2) by aligning histological sections to one another with rigid transformations (Supplementary Information A.2). In this step, all histological sections are used together, regardless of the type of biological information they represent. This initial histological volume is imperfect as it will contain sections that are not perfectly aligned to one another and also does not attempt to correct for 3D deformation artefacts in the sections. Despite the flaws in the initial reconstruction, it sufficiently recovers the true 3D anatomy of the donor brain such that it is possible to align to the reference structural volume, e.g., binary GM volume derived from T1w MRI of donor brain.

The goal of the next processing step is to align the histological sections to a structural reference brain volume (Figure 2.3) to correct the 3D deformations of the sections that occurred prior to the shock freezing of the tissue slabs described in Section 4.1. The alignment between the initial histological volume and the reference structural volume is done using an iterative strategy that proceeds from coarse to finer spatial resolutions (Supplementary Information A.3). The objective of this strategy is to produce a histological volume that more closely resembles the reference structural volume at each iteration. For the human brain, the reference volume was the post-mortem T1w MRI acquired from the donor. For the macaque brain the MEBRAIN template was used as a reference brain volume.

**Figure 3:**
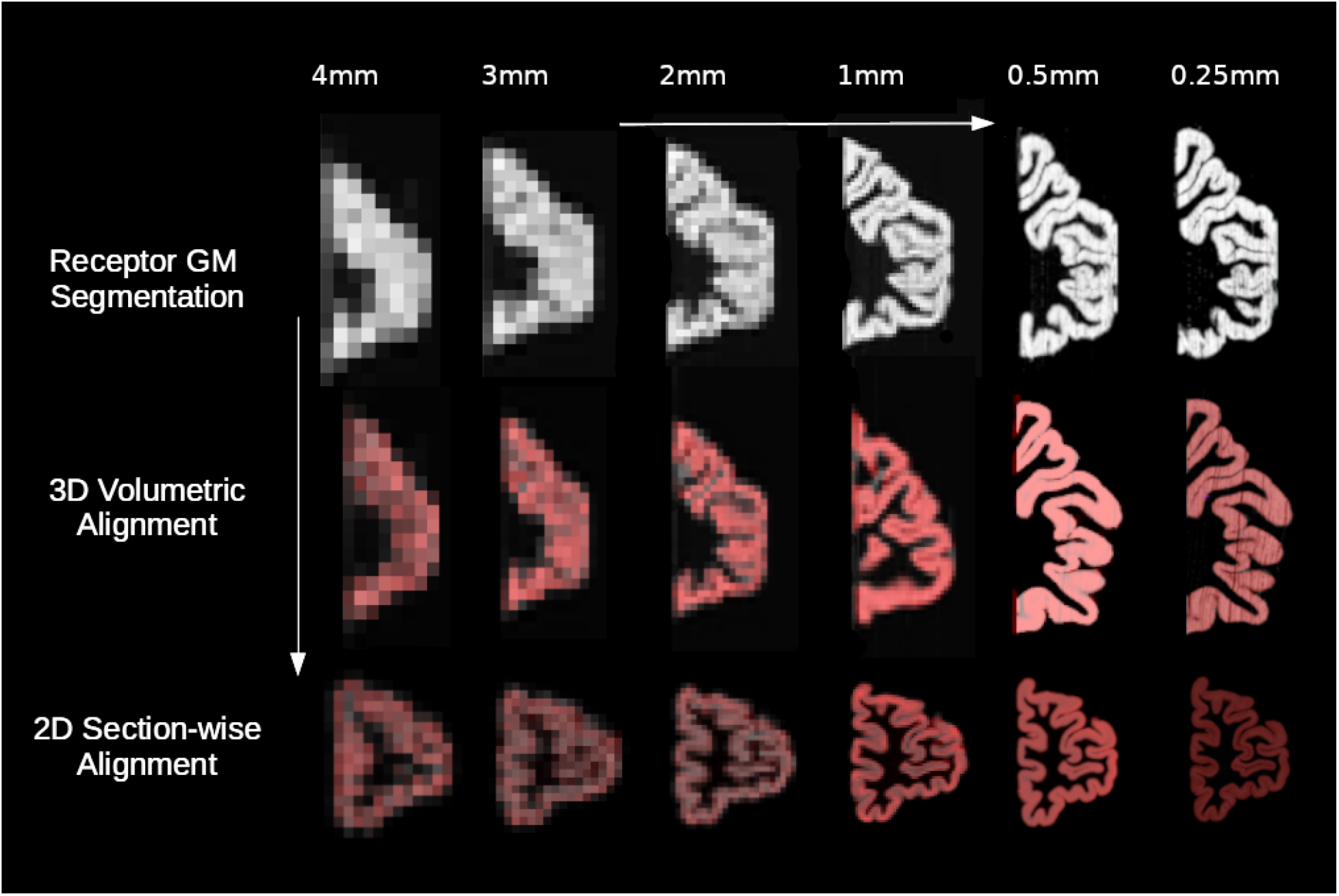
A multi-resolution scheme was used to align the histological slab volumes to the reference structural volume. At a given resolution, starting at 4mm, the histological sections are segmented into binary GM volumes. The binary histological GM volumes are non-linearly aligned in 3D to the reference structural volume. The individual histological sections are then aligned in 2D to the corresponding sections in the aligned reference structural volume.

**Figure 4:**
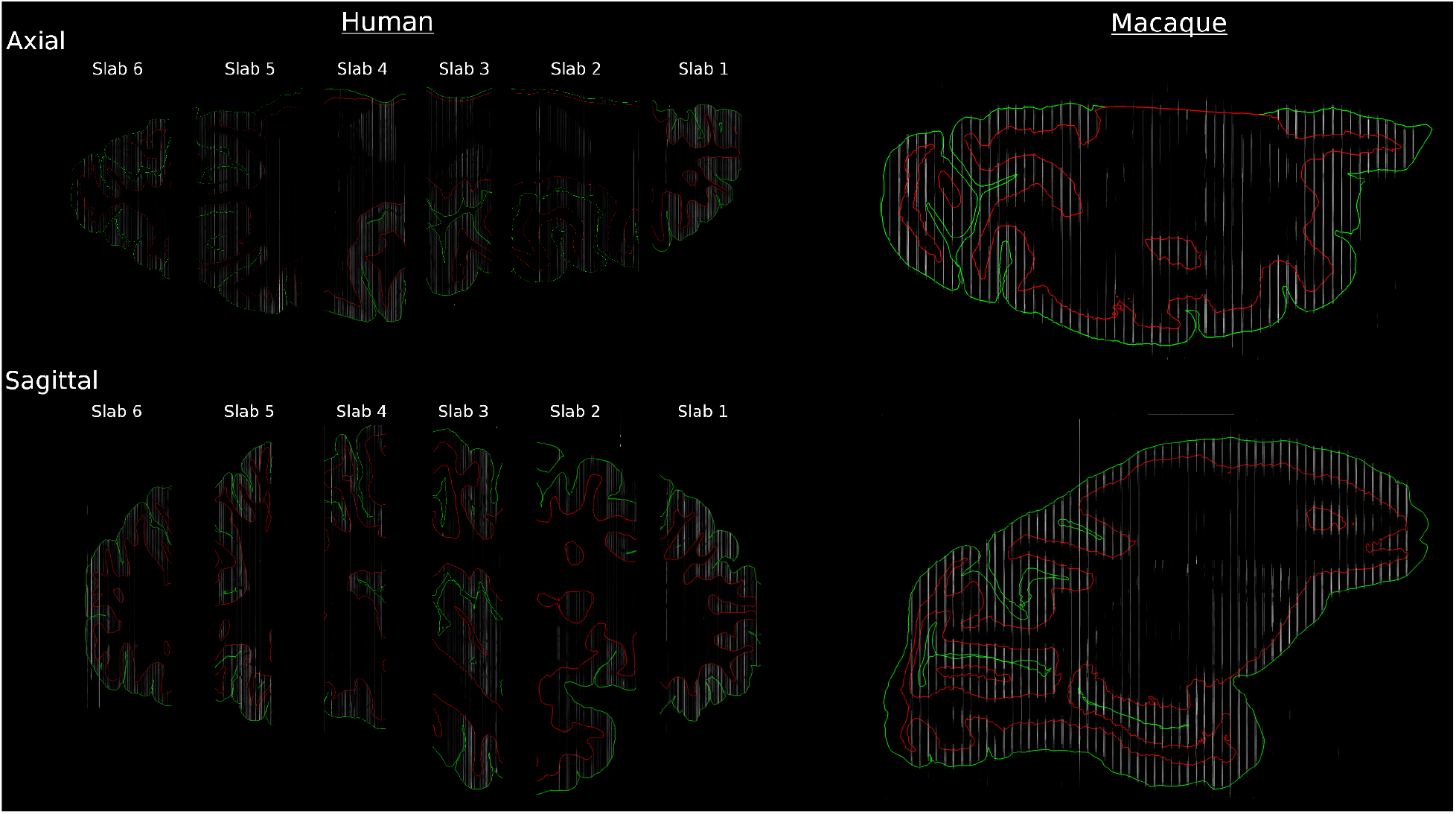
GM (green) and WM (red) cortical surface meshes superimposed on the 3D human (left) and macaque right) autoradiograph reconstructions. 3D volumes are shown in the receptor coordinate space of each tissue slab. Autoradiograph sections are almost all correctly aligned to the cortical surfaces.

**Figure 5:**
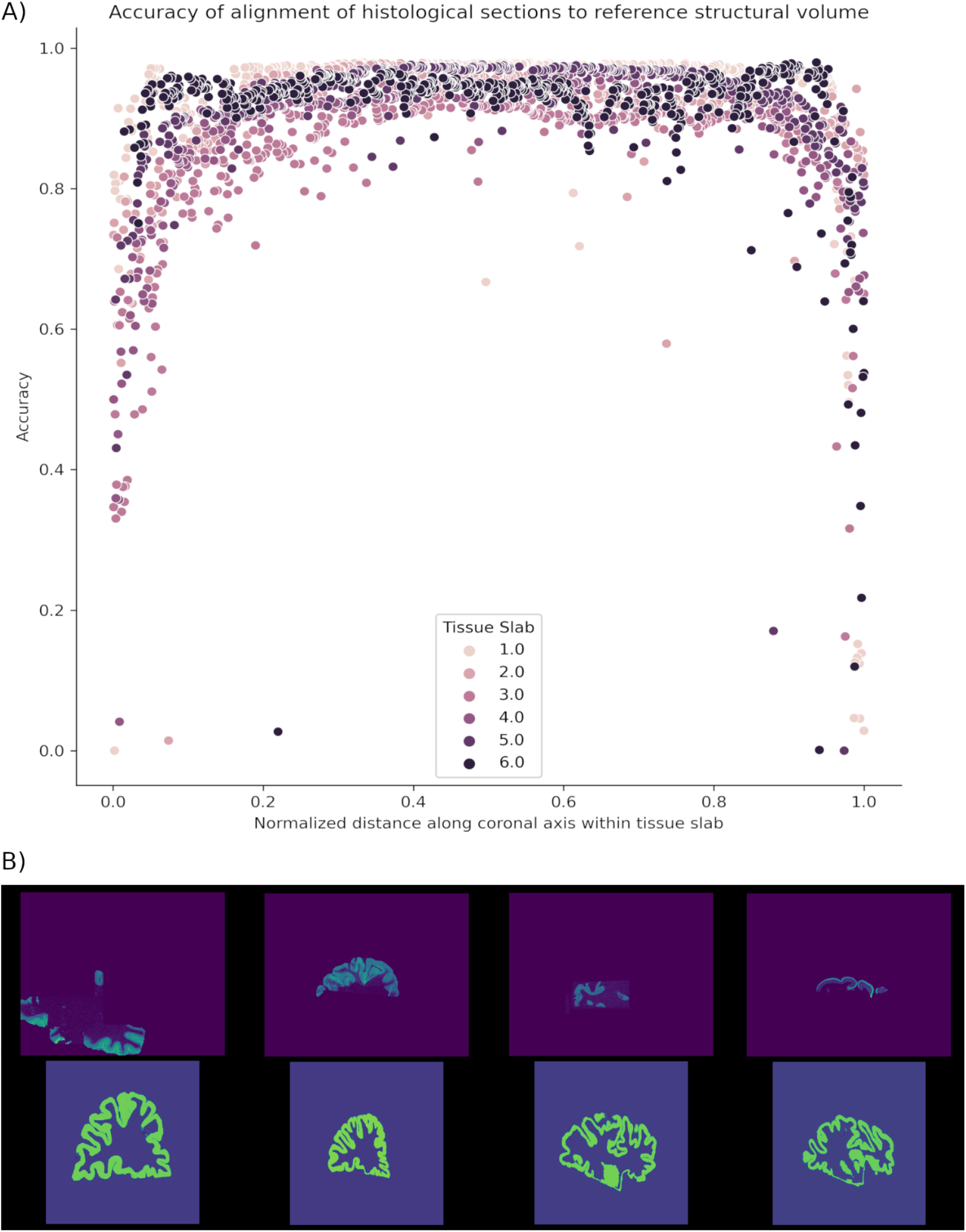
A) The accuracy (measured as the Dice score) of the alignment between thehistological sections and the reference volume MRI was plotted versus the relative position of the section within the tissue slab. Accuracy was significantly lower at the ends of the tissue slabs. B) 4 examples of histological sections from the ends of the tissue slabs (top row) illustrate the challenge of accurately aligning these sections to the corresponding sections from the reference structural volumes (bottom row).

**Figure 6.**
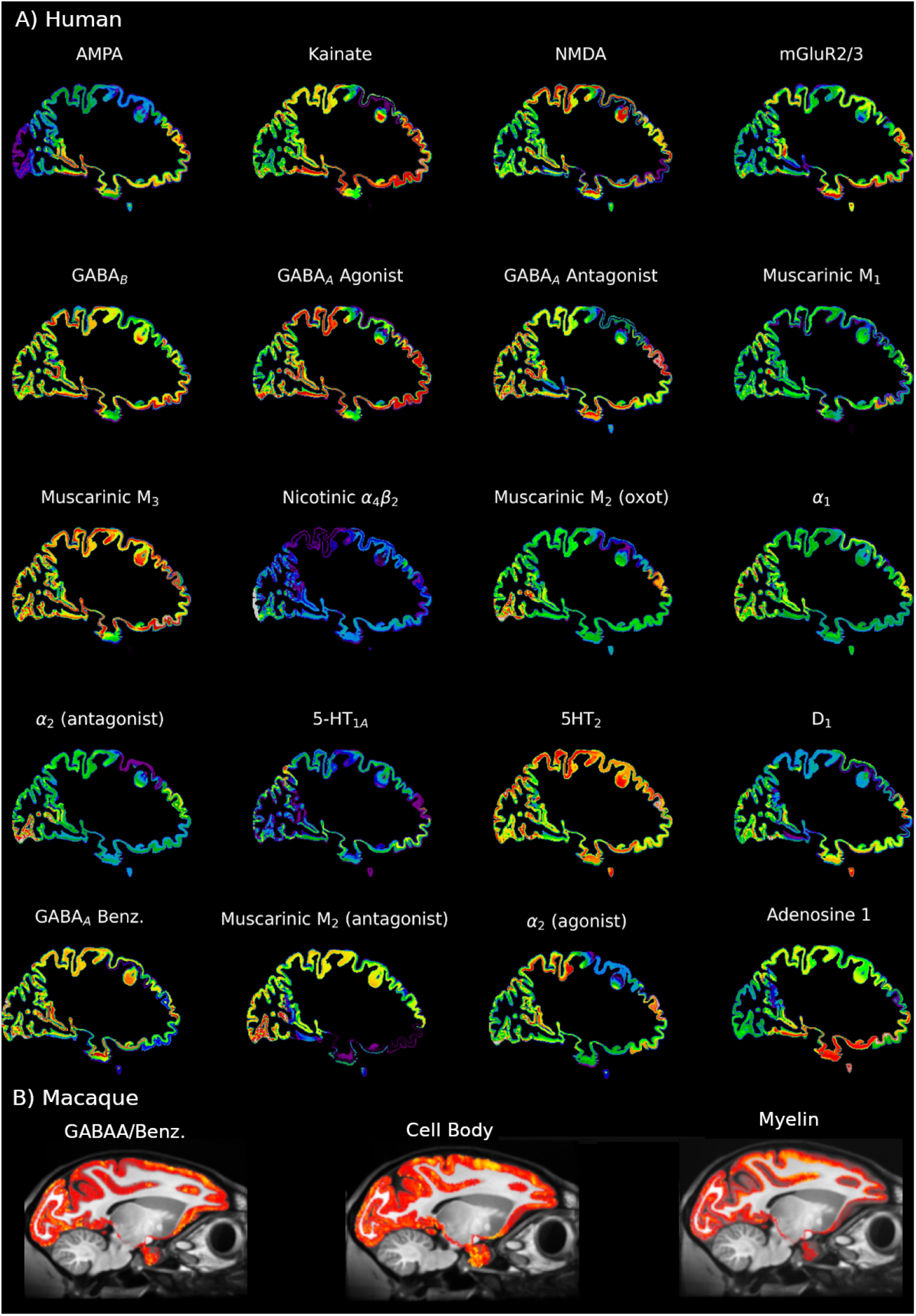
A) Sagittal sections resliced from the native coronal sections for each of the 20 neurotransmitter receptor binding sites for a human brain. Missing receptor binding densities are estimated using surface-based linear interpolation. B) Sagittal of the reconstructed volume of GABA_A_/BZ receptor binding density, the volume of reconstructed cell body sections, the volume of reconstructed myelin sections. The volumes were reconstructed using the MEBRAINS stereotaxic template because no MRI was acquired for the donor brain. Cell and myelin stained sections were not corrected for section inhomogeneity.

**Figure 7.**
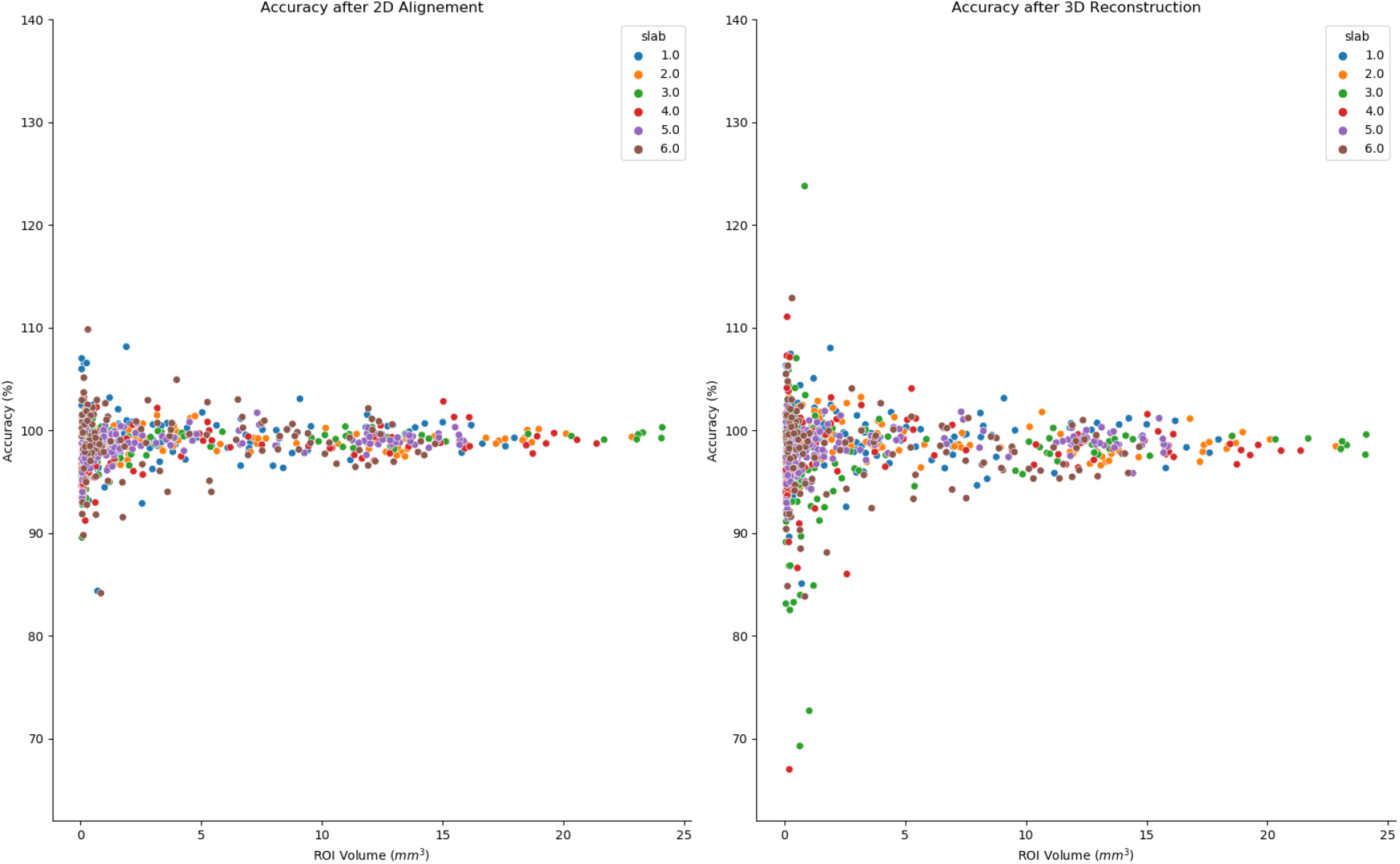
Binding densities were measured from human autoradiographs after applying 2D (left) and then 3D (right) nonlinear transformations to the autoradiographs and compared to binding densities measured from the raw autoradiographs.

**Figure 8.**
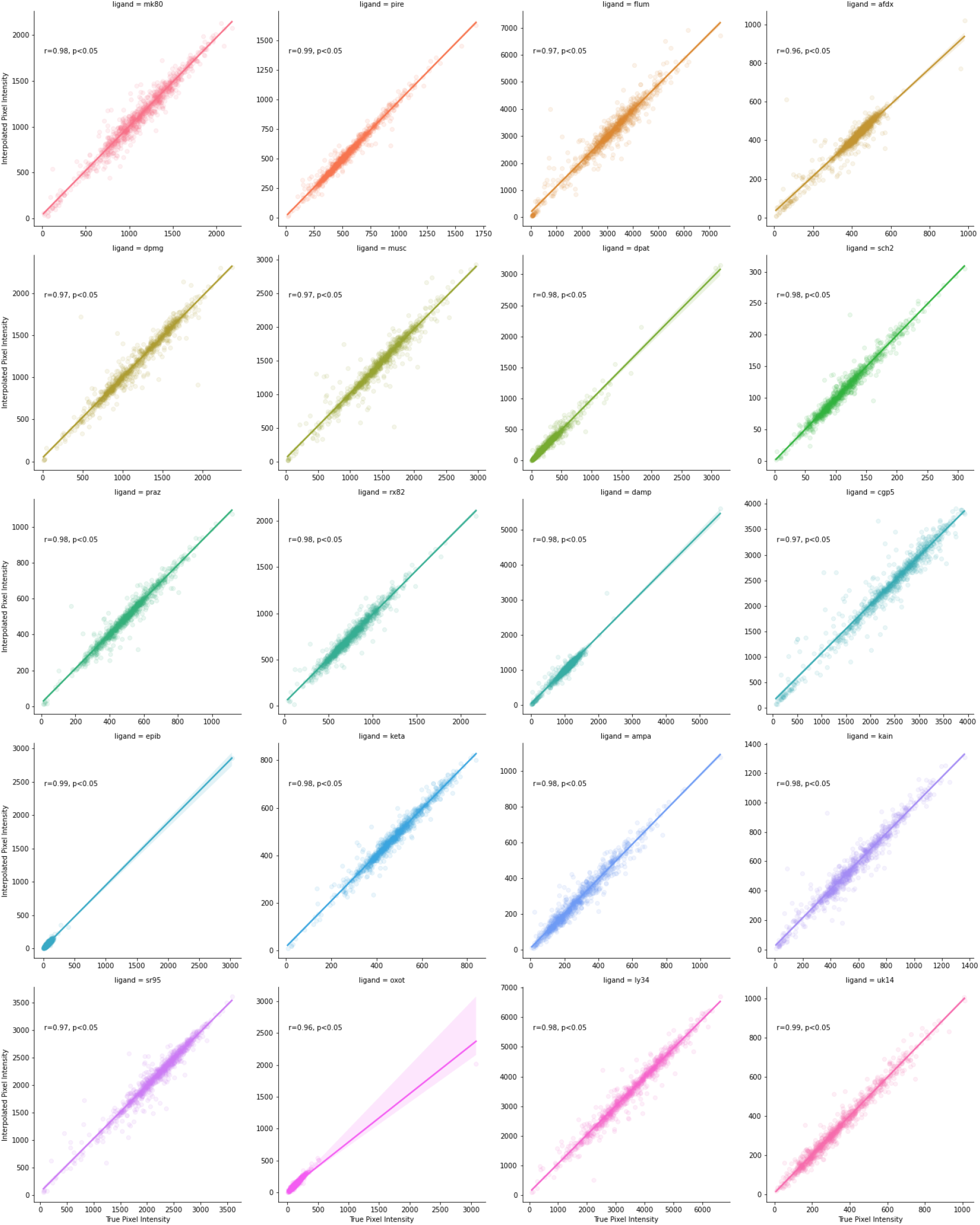
The surface-based interpolation algorithm was evaluated within acquired autoradiograph sections and demonstrated a high correlation between interpolated and true pixel intensities.

The processing steps for the multi-resolution alignment (Figure 2.3) are performed on each tissue slab independently, using an iterative, hierarchical series of progressively finer spatial resolutions. At each resolution the following steps are performed (Figure 3): a) the histological volume is segmented into a binary GM volume using a deep neural network trained for this task^38^; b) the histological GM volume is then nonlinearly aligned in 3D to the reference structural brain volume; and, finally, c) the 2D sections from the histological GM volume are non-linearly aligned in 2D to corresponding sections in the reference structural volume. Once completed at a given resolution, the processing steps are repeated at a higher spatial resolution until the processing is completed at the highest specified spatial resolution. The human data was reconstructed using a multi-resolution hierarchy of 4mm, 3mm, 2mm, 1mm, 500μm, 250μm to align the autoradiograph volume to the donor’s T1w MRI. The macaque reconstruction used the same hierarchy up to 1mm resolution.

Once the histological sections have been aligned to the structural reference volume up to the desired spatial resolution, the next task is to estimate the missing pixel intensities to create a reconstructed volumetric map for each type of histological section acquired (Figure 2.4). A surface-based method is used to interpolate missing pixel intensities for positions where no histological sections of a given type have been acquired (Supplementary Information A.4). For each type of section, there were large gaps between acquired sections and hence missing pixel intensities. Missing densities were estimated by projecting densities onto cortical surface meshes, interpolating missing densities over the cortical meshes, and interpolating surface receptor densities back into a volume.

### 2.3 Validation

#### 2.3.1 Validation of alignment of human and macaque brain sections

To demonstrate the efficacy of the reconstruction pipeline, it was applied to histological sections acquired from human and macaque brains, respectively.

The final reconstruction for the human data was generated at 250μm^3^. This resolution was chosen because it was sufficient to demonstrate the accuracy of the reconstruction and was the highest resolution feasible given available computational memory. All 20 available types of autoradiographs outlined in Table 1 were reconstructed.

The macaque data was only reconstructed up to 1mm as this was sufficient to demonstrate as a proof-of-principle that the pipeline can be applied a) to non-human data, b) using a reference template brain as reference structural volume, and c) using cell body and myelin stained sections in addition to autoradiography.

Quantitative validation of the alignment in the reconstruction was performed by calculating the Dice score between the aligned GM segmentation of the histological sections and the corresponding sections in the GM structural reference volume. The global accuracy of the reconstruction is calculated by taking the mean Dice score of each section, weighted by the total pixel intensity of the cortical GM in each respective section from the GM structural reference volume.

#### 2.3.2 Validation of the reconstructed pixel intensities

To ensure that the reconstruction pipeline did not affect the measured pixel intensities, regions in the unprocessed autoradiographs were compared to the corresponding regions in the reconstructed volumes. Regions of interest (ROIs) were generated based on similar patterns of pixel intensity distribution, ranging in area from 0.5mm^3^ to 24mm^3^. These ROI were not intended to be biologically meaningful but rather to appear similar to the real parcellation schemes, e.g., ROI from atlases such as the Julich Brain^41^, which we anticipate will be used to analyse the reconstructed data.

The parcellation of ROIs was produced by calculating a feature vector every 2mm on the raw autoradiographs (consisting of the mean, standard deviation, kurtosis, skew, and entropy of local pixel intensities). These features were then clustered using Gaussian Mixtures ^42^. All pixels within the brain were assigned the same label as that of the nearest clustered feature vector and all contiguous labelled regions were given a new, unique label value.

Finally, the autoradiograph parcellations were transformed using the same transformations as were applied to their corresponding autoradiographs. That is, a non-linear 2D transformation (A.3.3) and a 3D non-linear transformation (A.3.2). The accuracy of the transformed pixel intensities was quantified by calculating the correlation between true and interpolated pixel intensities.

As all histological sections are treated in the same manner, and thus are subject to the same potential interpolation errors, only sections from a single type of acquisition, autoradiography acquired with the [^3^H]-flumazenil ligand, were used to validate the reconstructed pixel intensities.

#### 2.3.3 Validation of surface-based interpolation

To validate the accuracy of the surface-based interpolation algorithm, we applied the surface-based interpolation algorithm within acquired autoradiograph sections For each ligand, respectively, 1,000 vertices that intersected acquired autoradiographs were selected at random. For each randomly selected vertex, the true pixel intensities were estimated based on the pixel intensities from neighbouring vertices. For each ligand, the Pearson r^2^ values were calculated between the true and estimated average pixel intensities in the core vertices (see Supplementary Information A.4.5).

## Code & Data Availability

The code used in this manuscript is available at www.github.com/tfunck/juelich-receptor-atlas. Reconstructed volumes for all brain and all types of histological sections will be made available to the public upon completion of the reconstruction of all brain hemispheres.

## Supplementary Information

### A.1 Preprocessing of sections and MRI

#### A.1.1 Automated cropping to isolate target brain tissue

The raw digitised images of the histological sections contain up to 3 components: the target section of brain tissue, extraneous pieces of brain tissue, and non-tissue objects (see Figure 1.D). The first step is to remove the extraneous pieces of brain tissue and non-tissue objects in the images. This is accomplished using an automated algorithm or manual cropping with the GIMP software when the automated approach fails.

### A.1.2 Extracting surfaces and grey matter mask from the reference structural volume

For the human brain, a binary MRI grey matter (GM) volume was extracted from the donor’s T1w MRI using a mesh representation of the cortical surface (Figure 9). A GM volume was used because the biological information represented in the histological sections creates a sharp contrast in pixel intensities between the GM and the rest of the image. Hence the GM in the histological sections can be aligned to the GM in the structural reference volume.

**Figure 9:**
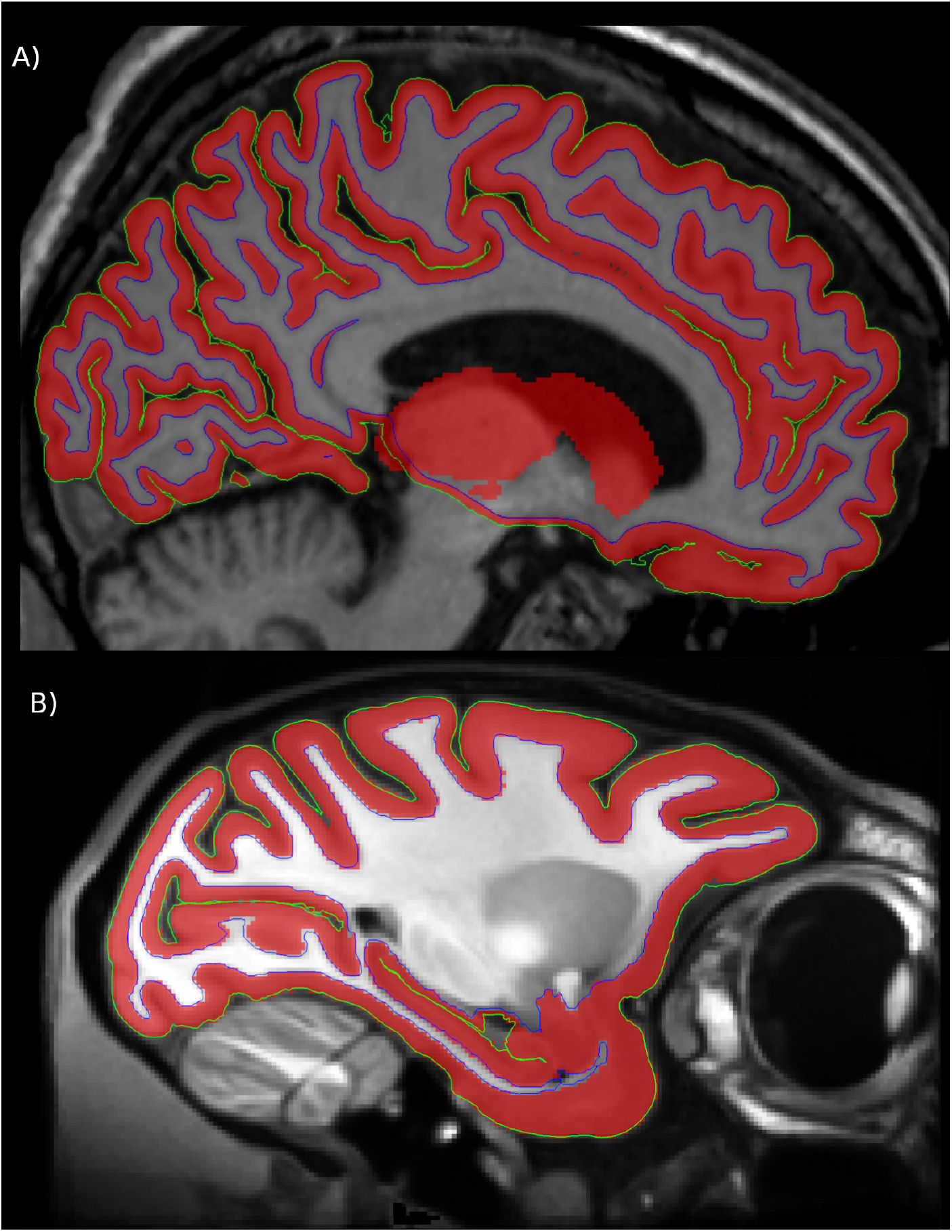
A) Cortical surfaces were extracted from the donor’s T1w MRI and used to derive a 250μm MRI GM volume. The ANIMAL algorithm was used to create a binary GM volume for subcortical GM regions. B) For the macaque reconstruction, Freesurfer was used to extract cortical surfaces and GM mask volume from the MEBRAIN template.

Cortical surface meshes were obtained from the MRI using the CIVET pipeline ^43^. A super-resolution cortical GM mask at 250μm was obtained from these cortical surface meshes by sampling points between the inner white-matter and outer GM surface meshes^44^. In principle, the reconstruction can be performed at an arbitrarily high resolution, but the presently available computational memory limited the reconstruction to 250μm. A segmentation of the donor’s subcortical GM was then generated using ANIMAL^45^, upsampled to 250μm using nearest neighbour interpolation, and added to the super-resolution cortical GM.

For the macaque reconstruction, Freesurfer^46,47^ was used to extract GM and white matter (WM) surfaces from the MEBRAINS template (Balan et al, 2022. In preparation). Hence for the macaque reconstruction, the reference structural volume did not come from the donor brain.

### A.2 Initial inter-section alignment

The initial step of 3D reconstruction is the alignment of the cropped histological sections to one another using 2D rigid body transformations. It is desirable to first align histological sections with high contrast before proceeding to lower contrast sections because low contrast sections are more likely to be misaligned and thereby propagate the misalignment to subsequent sections.

To this end, histological sections are first ranked automatically by the Michelson contrast ^48^ of pixel intensity produced by each method of histological acquisition. Histological sections produced with methods that yield images with the highest contrast are aligned first.

In each tissue slab, the most central histological section is considered as “fixed” in place and serves as the reference to which all subsequent sections will be aligned. Next, moving outwards from the central section of the slab, histological sections of the same type are aligned to their nearest “fixed” neighbour towards the center of the slab. Once a section has been aligned to its nearest neighbour, it is then considered “fixed” in place.

After the histological sections with the highest contrast have been aligned, they are all considered as “fixed” in place and can serve as a reference against which subsequent, lower contrast sections may be aligned. The alignment process therefore repeats with the histological sections with the next highest contrast and so forth until all sections have been aligned.

The rigid alignment is performed hierarchically with ANTs at voxel sizes of 4.096mm, 1.024mm, and 0.256mm with smoothing kernels of 2mm, 0.5mm, and 0.125mm full width at half-maximum (FWHM), respectively, and 100, 50, and 25 iterations ^49^.

For the reconstruction of the human brain, 6 initial histological volumes were produced by rigid alignment of the histological section, 1 for each tissue slab. For the macaque brain, the histological sections were treated as belonging to only a single slab and hence a single initial histological volume was produced for the acquired hemisphere.

### A.3 Alignment of histological volume to reference structural volume

#### A.3.1 Extracting GM mask from histological volume

After automated cropping of the histological sections, binary GM sections are generated from the cropped histological sections using a deep neural network trained to segment brain GM from 2D images^38^. Then at each resolution in the multiresolution hierarchy, these 2D histological GM sections are transformed using the best available transformations to align them together into a single 3D histological GM volume. For the first resolution in the multiresolution hierarchy, the best available transformations are the rigid body transformations calculated in the initial inter-section alignment (Supplementary Information A.1). At subsequent resolutions, the best alignments are the nonlinear 2D transformations calculated in the previous resolution in the hierarchy (A.3.3). Therefore, after the first step of the multiresolution hierarchy, the 2D histological GM volumes are transformed so that they better correspond to the actual anatomy of the reference structural volume to which they were aligned at the previous resolution of the hierarchy.

The aligned 2D histological GM images contain gaps along the coronal axis where no sections were acquired and which must be filled to produce a continuous representation of the cortical GM. Nearest neighbour interpolation between aligned 2D histological GM images is used to estimate the morphology of the cortex where no histological sections were acquired. The 3D histological GM volume is then downsampled to the current resolution in the resolution hierarchy with an order 5 spline interpolation. This produces the continuous 3D histological GM volumes illustrated in the top row of Figure 3.

#### A.2.3 3D alignment of histological volumes to reference structural volume

When more than one slab of tissue is reconstructed, as in the human data set, a significant challenge is to identify which portion of the reference structural volume corresponds to each slab of brain tissue. Due to the deformation of the tissue slabs prior to freezing and loss of sections between slabs, the total width of the brain slabs along the coronal axis was less than that of the brain in the MRI volume, hence the slabs could not simply be placed adjacent to one another. We manually identified the anterior and posterior most points of each slab on the reference structural volume and extracted a corresponding slab of tissue from the reference volume. Therefore the alignment of the receptor slab was limited to a manually defined portion of the corresponding reference structural volume.

For each slab, the GM volume for the sections is linearly aligned to the portion of the reference structural volume with rigid (3 rotations and translations), similarity (3 rotations and translations, 1 global scaling), and full affine transforms (3 rotations, translations, scaling, and shearing parameters). After the linear alignment, SyN is used to perform non-linear alignment between the slab GM volume and the corresponding portion of the reference structural volume. Alignment is performed with ANTs using 1000 iterations for each alignment with the Mattes mutual information metric^50^.

This step may be omitted if histological sections sampled from a single tissue slab spanning a whole hemisphere are being reconstructed, as was effectively the case for the macaque data set.

#### A.3.3. 2D refinement of histological section alignment to reference structural volume

After aligning the histological GM volume to the reference structural volume in 3D, the alignment is refined by aligning the histological sections in 2D to their corresponding coronal sections in the reference structural volume. This 2D alignment between corresponding coronal sections is possible because the 3D alignment in the previous stage produces a reference structural volume that has been transformed into the coordinate space of the histological GM volumes for each tissue slab. The alignment is performed with ANTs^49^ using rigid, similarity, affine, and nonlinear (SyN) 2D transformations using the Mattes mutual information^50^ similarity metric.

### A.4 Surface-based interpolation of missing binding densities

#### A.4.1 Upsampling surface meshes

Intermediate cortical surfaces are generated between the WM-GM and pial-GM border by evenly subdividing the depth between the 2 borders. The number of intermediate cortical surfaces is calculated such that each pixel in the acquired histological sections is intersected by at least one cortical surface. This is estimated by dividing the maximum cortical thickness of the brain being reconstructed by the final resolution of the reconstruction.

Each of the cortical meshes is then supersampled such that the maximum distance between any two neighbouring vertices is less than or equal to the final resolution of the reconstruction, i.e., 250μm for the human reconstruction and 1mm for the macaque reconstruction (Figure 10.C). This upsampling step is done to ensure that there is at least one vertex per voxel in sections where histological sections have been acquired.

**Figure 10.**
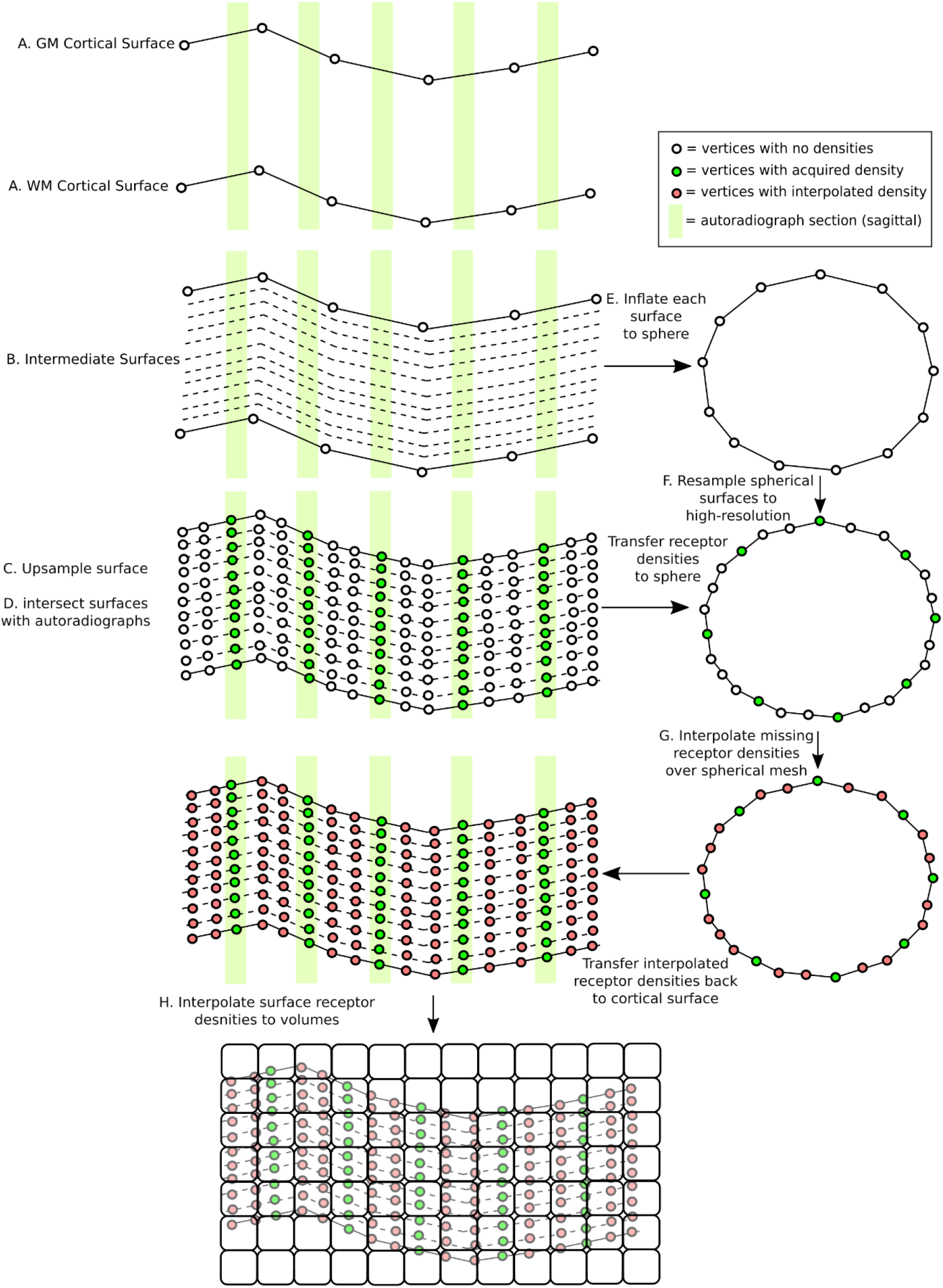
Illustrative schema of surface-based interpolation algorithm. Intermediate meshes are defined between the WM and GM surfaces. The cortical surfaces are upsampled and pixel intensities of autoradiographs are projected onto the surface (green circles). Each of the surfaces is inflated to a sphere. Missing pixel intensities are interpolated over the spherical surfaces (red circles). The acquired and interpolated pixel intensities are interpolated into a volume to produce a volumetric atlas of receptor density for a specific receptor.

For the human brain, 28 intermediate cortical surfaces were generated, yielding a total of 30 cortical meshes spanning the depth of the cortex between the WM-GM and GM-pial border (Figure 10.B). In the macaque brain, 8 intermediate cortical surface meshes are generated between the WM-GM and GM-pial border, yielding a total of 10 cortical surface meshes. The number of surfaces would be increased for higher resolution reconstructions such that at least one intermediate surface would intersect every pixel between the WM-GM and GM-pial border.

#### A.4.2 Projecting binding densities onto a cortical surface mesh

To minimize potential interpolation artefacts on the histological sections, the surface meshes are transformed from the coordinate space of the reference structural volume to the coordinate space of each of the slab volumes, respectively, by applying the inverse linear transformations and non-linear deformation fields calculated in section (Supplementary Information A.3.2). The pixel intensities in histological sections are projected onto the surfaces in the native coordinate space of the slab.

The histological sections are resampled to the final resolution of the reconstruction and interpolated, with nearest neighbours, to the transformed and upsampled cortical mesh for that slab (Figure 10.D).

#### A.4.3 Interpolating missing binding densities

After interpolating pixel intensity values from histological sections to the transformed cortical surface meshes, missing vertex values are estimated for vertices at which no pixel intensities were measured. All of the meshes along the depth of the native cortical surfaces are inflated to spheres (Figure 10.E) using the Freesurfer’s mris_sphere (iterations = 500)^46^. The inflated spherical meshes are then resampled to the same number of vertices as the upsampled surfaces (Figure 10.F). Therefore, for each vertex in the upsampled cortical mesh, there is a corresponding vertex on the upsampled inflated sphere.

The missing pixel intensities for each cortical surface mesh are interpolated by applying linear interpolation over each of the corresponding upsampled inflated spherical meshes (Figure 10.G) using the SSRFPACK algorithm^51^ implemented in the stripy^52^ Python package. The interpolation was performed on the inflated sphere instead of directly on the cortical surfaces, because it is computationally simpler to calculate distances between vertices on a simple geometric object like a sphere than on the complex surface of the cortex. Furthermore, given that the surface inflation process approximately preserves relative distances between vertices^46^, the interpolated densities on the inflated sphere are equivalent to those which would have been calculated directly on the cortical surface.

#### A.4.4 Interpolating surface pixel intensities to a volume

Finally, a volumetric surface was generated from the surface pixel intensity values by interpolating voxel values from vertex values across the depth of the cortical surfaces (Figure 10.H). Specifically, the pixel intensities for voxels lying within the GM and WM surface meshes was set equal to the mean receptor density of any vertices contained within the volume of a voxel. This method may leave gaps in the reconstructed cortex where no surface vertices are located within the volume of a voxel. The voxel intensities of these empty voxels are estimated by linear interpolation based on the values of the neighbouring voxels with pixel intensities.

The surface-based interpolation was applied independently to each slab of reconstructed tissue for each type of acquired histological section to estimate missing pixel intensities *within* the tissue slab. The individual tissue slabs were then transformed into the coordinate space of the reference structural image. Finally, pixel intensities *between* the tissue slabs were estimated by again applying the surface to voxel interpolation described in this section for voxels located between the tissue slabs.

#### A.4.5 Validation of surface-based interpolation

For each ligand, respectively, 1,000 vertices that intersected acquired autoradiographs were selected at random. For each of these “seed” vertices, neighbours were identified within *n* steps along the surface mesh within the same plane as the acquired autoradiograph (all vertices in Figure 11), where *n* follows a uniform probability distribution *n ~ U(2,6)*. For the purpose of validation, the vertices from the seed vertex to the n-1 neighbour (purple, green, and blue vertices in Figure 11) were treated as though the pixel intensities at these locations were missing, though in fact all pixel intensities were known because the vertices all intersect an acquired autoradiograph. A subset of *m* neighbours around the seed vertex, where *m ~ U(1,5)*, were then identified (purple and green vertices in Figure 11). These form a core patch of vertices whose average pixel intensity was estimated.

**Figure 11.**
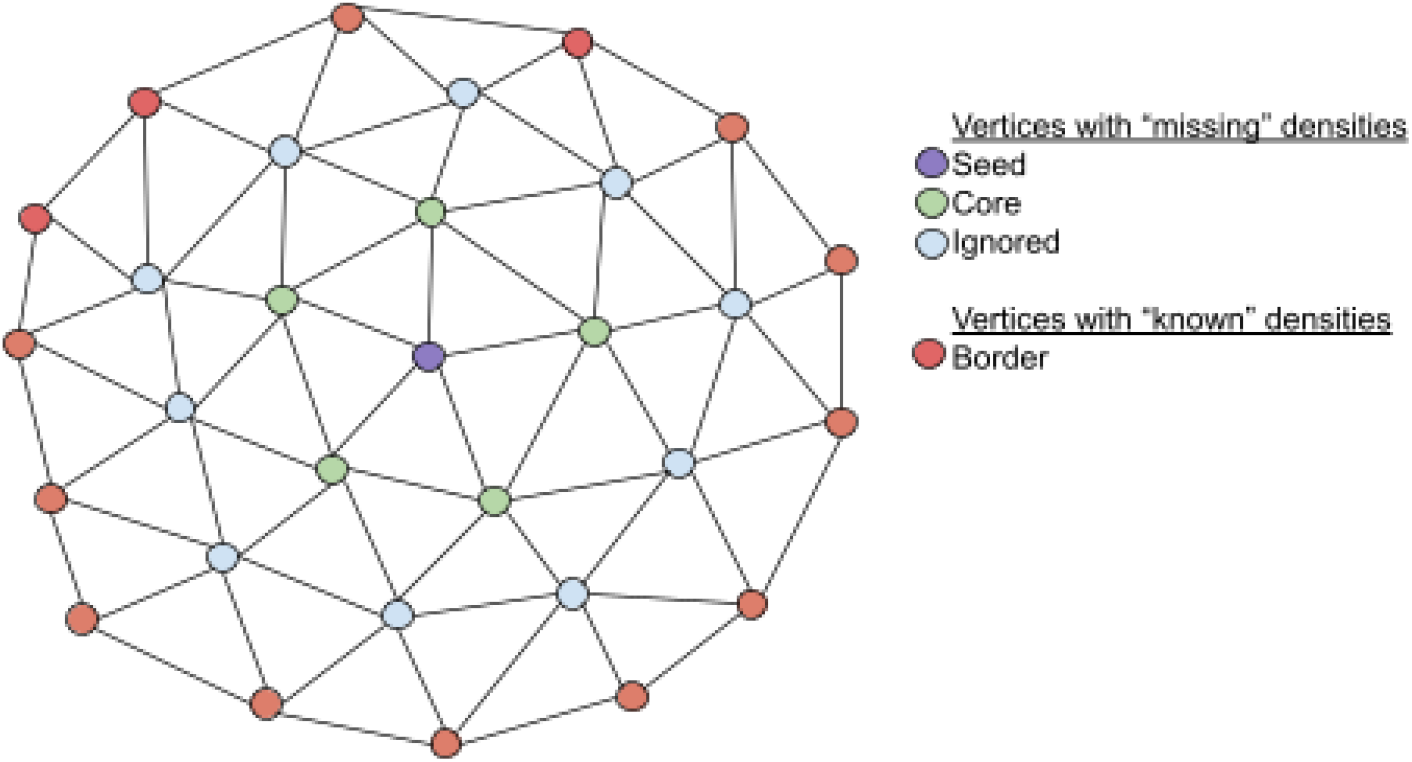
A toy example of a mesh patch used to validate the surface-based linear interpolation for estimating missing pixel intensities. Here a seed vertex, purple, with a neighbourhood of core vertices, *m=1*, is estimated using known border vertices, red, *n=3*.

The vertices that were *n* edges away from the seed vertex (red vertices in Figure 11) were considered to have known pixel intensities. The surface-based linear interpolation algorithm was then used to estimate pixel intensities for vertices within the seed and core patch of vertices given the vertices with “known” intensities. Pixel intensities were estimated for each vertex individually and then averaged together.

